# Transformer-based deep learning integrates multi-omic data with cancer pathways

**DOI:** 10.1101/2022.10.27.514141

**Authors:** Zhaoxiang Cai, Rebecca C. Poulos, Adel Aref, Phillip J. Robinson, Roger R. Reddel, Qing Zhong

## Abstract

Multi-omic data analysis incorporating machine learning has the potential to significantly improve cancer diagnosis and prognosis. Traditional machine learning methods are usually limited to omic measurements, omitting existing domain knowledge, such as the biological networks that link molecular entities in various omic data types. Here we develop a Transformer-based explainable deep learning model, DeePathNet, which integrates cancer-specific pathway information into multi-omic data analysis. Using a variety of big datasets, including ProCan-DepMapSanger, CCLE, and TCGA, we demonstrate and validate that DeePathNet outperforms traditional methods for predicting drug response and classifying cancer type and subtype. Combining biomedical knowledge and state-of-the-art deep learning methods, DeePathNet enables biomarker discovery at the pathway level, maximizing the power of data-driven approaches to cancer research. DeePathNet is available on GitHub at https://github.com/CMRI-ProCan/DeePathNet.

**Highlights:**

- DeePathNet integrates biological pathways for enhanced cancer analysis.
- DeePathNet utilizes Transformer-based deep learning for superior accuracy.
- DeePathNet outperforms existing models in drug response prediction.
- DeePathNet enables pathway-level biomarker discovery in cancer research.

## Introduction

Multi-omic analysis of diverse data types enables researchers to gain insights into tumor biology and identify new and robust therapeutic targets [1]. One major goal of multi-omic analysis by machine learning is to predict the cancer treatment strategies that are best suited to individuals in the context of precision medicine. A variety of multi-omic studies have led to the improved detection of intra-tumor heterogeneity, identification of novel therapeutic targets, as well as more robust diagnostic and predictive markers [2–5]. Many of these discoveries would not have been possible by analyzing any single omic data type alone. However, performing multi-omic analysis presents computational challenges due to the large amounts of data generated by high-throughput instruments and the limitations of existing multi-omic data integrative methods [6,7].

To address this, a plethora of machine learning methods have been developed for integrating large-scale multi-omic data [2–4,7–11]. For example, moCluster [8] integrates multi-omic data based on joint latent variable models, showing performance superior to previous methods such as iCluster [9] and iCluster Bayes [10]. Likewise, mixOmics [11] provides various options for multi-omic data integration, aiming to find common information among different omic data types. These models solely take omic measurements as the input and do not consider existing biomedical knowledge that links different omic data types together, such as regulatory networks. Regulatory networks exist in cells to control the expression levels of various gene products through collections of functionally interacting protein or RNA macromolecules [12]. However, models that incorporate existing biomedical knowledge in addition to computational inference have the potential to better capture the interactions that drive biomarker associations, and to increase the predictive power and modeling capacity of these algorithms [7].

In omic studies, most existing attempts to incorporate biomedical knowledge into machine learning are limited to one single omic data type [13–16]. ATHENA [17] and PARADIGM [18] support multi-omic data, but they are based on linear models that are not complex enough to model the relationships among pathways. Using deep learning, several studies have attempted to incorporate existing biomedical knowledge into multi-omic models [3,19]. DCell [20] and DrugCell [21] combine the neural network architecture with known gene ontology information, but they only support using gene deletions or mutation as the input. EMOGI [22] was designed based on graph neural networks (GNN) [23,24] and it integrates protein-protein interaction (PPI) networks with multi-omic data to predict cancer genes. However, its network architecture cannot be easily generalized to other tasks. Furthermore, a GNN-based model was developed to classify breast cancer subtypes using multi-omic data, but the model is not interpretable for discovering potential new biological mechanisms or drug targets [25]. Besides, gene ontology information and PPI networks used in these models do not precisely reflect cancer-specific information. Therefore, integrating cancer pathways [26] into multi-omic data analysis by deep learning for general tasks, such as drug response prediction and cancer type or subtype classification, remains an open research topic.

Inspired by the computational foundation of Generative Pre-trained Transformer (GPT) [27] that significantly contributed to the recent achievements of ChatGPT [28], we developed DeePathNet, a Transformer-based [29] explainable deep learning method that integrates multi-omic data, such as genomic mutation, copy number variation, gene expression, DNA methylation, protein intensity, and CRISPR-Cas9 data, among others, with knowledge of cancer pathways. The Transformer architecture has enabled many breakthroughs in artificial intelligence [30–32]. In molecular biology, the Transformer has been used to model DNA sequence, protein tertiary structures, and drug chemical structure data[33–35]. Furthermore, the Transformer-based models have been used together with GNN to analyze mRNA and miRNA data in cancer, but these models are not interpretable [36,37]. Therefore, whether the Transformer can be combined with cancer pathway information to integrate any number of omic data layers for multi-omic cancer data analysis is unknown.

Our contributions in this paper can be summarized as follows:

1. A novel deep learning model called DeePathNet has been developed for analyzing cancer molecular data. The novelty of DeePathNet lies in its unique design, which enables the Transformer to solve previously unanswered questions in cancer research. This offers a fresh avenue for employing Transformer architecture in the analysis of multi-omic data within the field of cancer. By combining the Transformer with domain-specific knowledge of cancer pathways, DeePathNet not only surpasses the predictive accuracy of conventional machine learning approaches but also provides reliable model explanations.
2. The performance of DeePathNet has been evaluated and validated using large-scale datasets and multiple evaluation metrics, which are not commonly included in similar studies. Exemplary tasks, including drug response prediction and cancer type and subtype classification, have been selected due to the availability of relatively large amounts of data. DeePathNet can be extended to other tasks as well.
3. The top omic features and cancer pathways have been reported using DeePathNet’s feature importance, facilitating the discovery of potential new biomarkers. This information provides valuable insights into the underlying mechanisms of cancer development and progression.

## Methods

### Multi-omic and drug response data collection

For drug response prediction, multi-omic data were retrieved from 941 CLP [38] and 696 CCLE cell lines [39]. In total, 19,099 gene mutations, 19,116 CNV and 15,320 gene expression features are in the CLP, and 18,103 gene mutations, 27,562 CNV, and 19,177 gene expression features are in the CCLE.

For drug response prediction analysis with proteomic data, the ProCan-DepMapSanger dataset [40] was added to the CLP (CLP + ProCan-DepMapSanger = CLP^+^) and the CCLE’s proteomic dataset [41] was also used (CCLE + CCLE proteomic data = CCLE^+^). The ProCan-DepMapSanger and CCLE proteomic datasets contain 8,498 and 12,755 protein features, respectively. The combined datasets have 910 and 292 cell lines for CLP^+^ and CCLE^+^, respectively. No additional processing was performed on the datasets (Table S1). When using the CCLE and CCLE^+^ as the independent test set, we excluded the cell lines that are also in the CLP dataset, and only used the 71 and 33 cell lines that are unique in the CCLE and CCLE^+^ datasets, respectively.

For cancer type and subtype classification, multi-omic data from TCGA cohorts were retrieved using TCGA-assembler 2 [42]. In total, 6,356 samples were collected, containing 31,949 features from gene mutation, 23,529 features from CNV and 20,435 features from gene expression. In addition, multi-omic data from 122 breast cancer samples were retrieved from a CPTAC breast cancer cohort [43], containing 11,877 features from gene mutation, 23,692 features from CNV and 23,121 features from gene expression. For breast cancer subtype classification, the PAM50 classification (Luminal A, Luminal B, HER2+, Basal and Normal-like) was retrieved from the TCGA and CPTAC datasets (Table S1).

### Overview of DeePathNet

DeePathNet was developed to model biological pathways using a Transformer-based deep learning architecture with both multi-omic data and cancer pathway information as the input (Figure 1A). The performance of DeePathNet was evaluated on drug response prediction, and cancer type and subtype classification.

**Figure 1.**
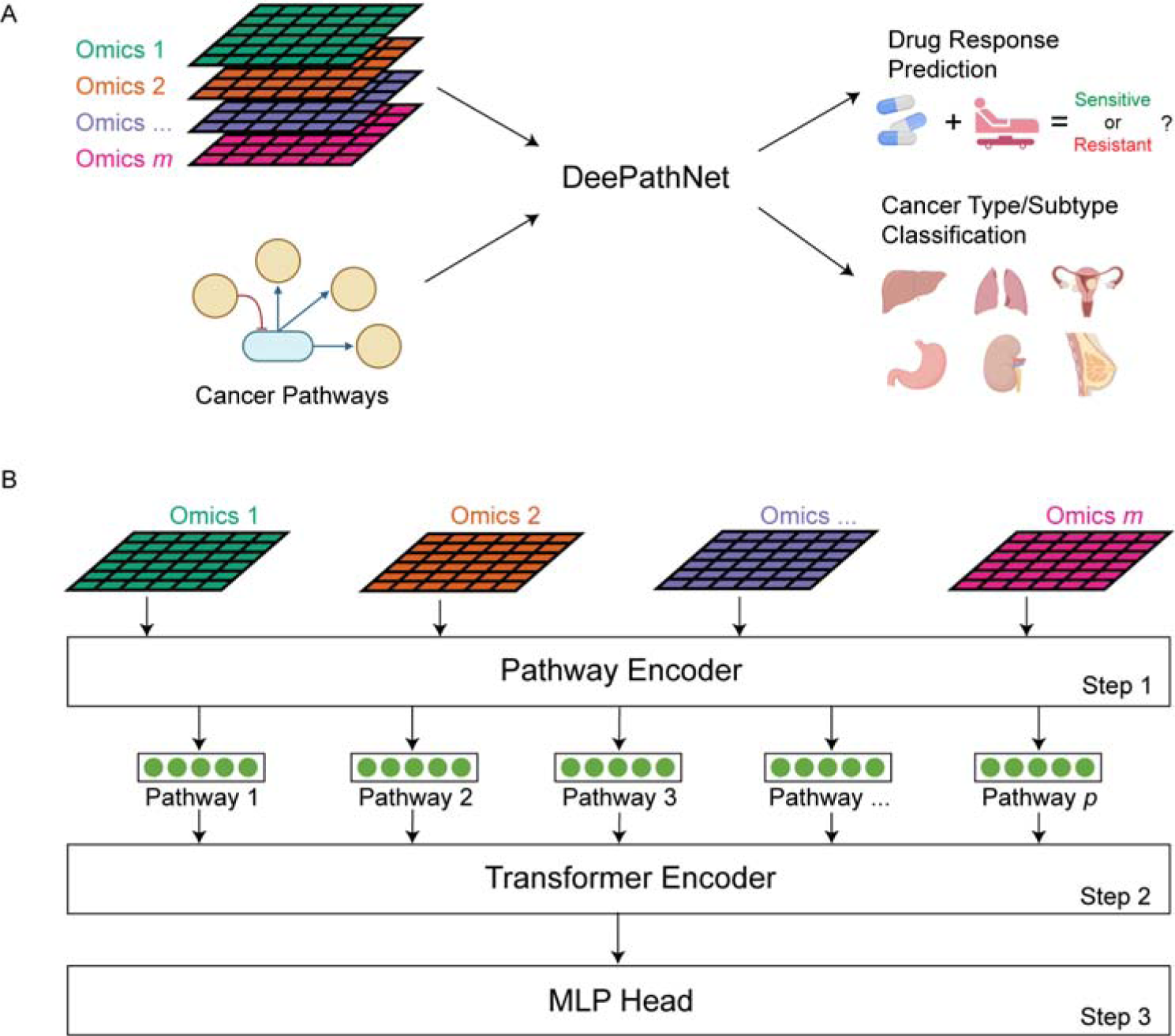
Overview of DeePathNet. **A,** DeePathNet has its network architecture built using the LCPathway dataset and takes multi-omic data as the input to model pathway interactions and predict drug responses or classify cancer types and subtypes. **B,** DeePathNet architecture supports any number of omic data types as the input. Step 1: DeePathNet encodes multi-omic information into cancer pathways. Step 2: DeePathNet uses a Transformer encoder to learn the interactions among these pathways. Step 3: The encoded pathway vector is passed into an MLP for the prediction. Circles represent neurons in a neural network. Arrows represent the direction of information flow.

DeePathNet consists of three major steps. It starts with a pathway encoder to summarize features from omic data types into cancer pathways (Step 1; Figure 1B), and then uses a Transformer encoder to model the interactions among these pathways (Step 2). This is followed by a multi-layer perceptron (MLP) that can be adapted to different prediction tasks (Step 3).

In Step 1, the neural network architecture is constructed based on the LCPathways dataset,[26] which contains 241 literature-curated pathways encompassing 3,164 cancer genes. The LCPathways dataset was selected since it is one of the most recent and comprehensive pathway databases that are specifically curated for cancer research. As such, it is particularly suitable for the applications of DeePathNet. The pathway encoder then uses a fully connected layer to project the multi-omic data (Omics 1–*m*) from genes (Gene 1–*n*) onto a 512-dimension pathway vector that represents one of the cancer pathways (Pathway 1– *p*; Figure S1A, see Experimental Procedures). With this architecture, the pathway encoder allows DeePathNet to capture interactions across different omic data types. Moreover, grouping multi-omic features into pathways reduces the dimensionality of the original high-dimensional data. This mitigates the challenge of the curse of dimensionality, making it computationally feasible to apply the model to large datasets without sacrificing its performance [44].

In Step 2, an enhanced version of the Transformer module is developed to encode the interactions among cancer pathways (Figure S1B, see Experimental Procedures). First, a dropout layer is used to train only half of the pathways at each iteration, which prevents the model from focusing on specific pathways that may not generalize well to a test dataset. Then, two blocks of the original Transformer module [29] are used, which contains a list of recurring layers, with each layer comprising a sequence of layer normalization, multi-head self-attention, and an MLP. The Transformer also enables dynamic modeling of the complex relationships among cancer pathways, thus avoiding the generation of fixed weights for the different inputs, as in traditional machine learning.

In Step 3, an MLP is used to map the encoded pathway vectors to output neurons, which allows the knowledge learned by the Transformer module to be adapted to general prediction tasks. The three components of DeePathNet work together and cannot be applied to multi-omic data individually.

### Details of DeePathNet

DeePathNet has a pathway encoder (Step 1), a Transformer encoder (Step 2) and a MLP (Step 3).

In Step 1, DeePathNet encodes multi-omic information into cancer pathways, defined by the 241 cancer pathways in LCpathways [26]. Let *g_mutation_* ∊ {0,1} represent the mutation, *g_CNV_* ∊ ℝ the CNV, *g_RNA_* ∊ ℝ the gene expression, and *g_pro_* ∊ ℝ the protein intensity of a gene *g*. Then the vector that contains omic features for a pathway that contains *n* genes with four omic data types, is defined as:

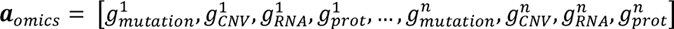

Next, the vector ***a**_omics_* is encoded into the pathway vector ***a**_encoded_* via an MLP. Here, the notation is converted into the matrix form to include the number of samples. Thus, for *N* samples, the total features from the four omic data types for a pathway can be represented as a matrix ***A**_omics_* of dimension *N* × 4*n*. DeePathNet then uses a fully connected layer to encode these omic features into an encoded pathway matrix ***A**_encoded_*, calculated as:

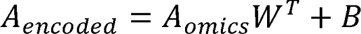

where *W* and *B* represent the learnable weights matrix and bias term in the fully connected layer. The dimension of the weight matrix *W* is set as 512 x 4*n* and the dimension of both *A_encoded_* and *B* is *N* x 512. In total, 241 cancer pathways were used, and 241 matrices 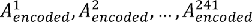 are combined as a tensor ***A****_encoded_* with a dimensionality of *N* × 512 × 241. ***A****_encoded_* is used as the input into the Transformer encoder (Figure S1).

To assess the predictive power conferred by the LCPathway dataset, we configured DeePathNet such that genes within each pathway were replaced with randomly selected genes, while maintaining the original size of each pathway. When DeePathNet was trained on the CLP dataset and subsequently evaluated on the CCLE dataset for drug response prediction, we observed superior predictive performance using LCPathways compared to pathways populated with randomly selected genes for the majority of the evaluation metrics (Figure S6A). This disparity in performance became more pronounced when the model was applied to the breast cancer subtype classification task. Specifically, when trained on the TCGA dataset and tested on the CPTAC dataset, the use of LCPathways resulted in a marked enhancement in predictive performance across all three evaluation metrics under consideration (Figure S6B). These results substantiate the notion that LCPathways contributes meaningful biological information, thereby augmenting DeePathNet’s predictive capabilities.

In Step 2, DeePathNet uses a Transformer encoder to learn the interdependence among regulatory pathways in cancer. In contrast to the general attention mechanism that models the interdependence between the input and target, self-attention is used by the Transformer module to model interdependence within the input [45] (i.e., features from the multi-omic data). The Transformer encoder starts with a dropout layer with a probability of 0.5 on the 241 cancer pathways, ensuring that on average half of the pathways are dropped out during training to prevent potential overfitting. The set of selected pathways is sampled independently for each training batch, allowing different pathways to be used. The Transformer block was configured the same way as the original version [29], denoted as *Transformer* below. Since the Transformer encoder contains recurrent layers, we use a superscript with parenthesis to represent the ***A***_encoded_ at different layers, where 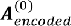 represents the data before entering the first layer. After the first layer of the Transformer block, 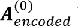 becomes 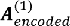 as follows:

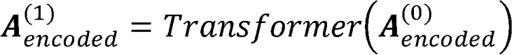

DeePathNet contains two layers of Transformer block, therefore:

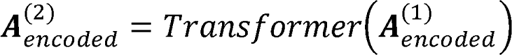

Finally, in Step 3, DeePathNet uses a MLP to map 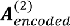 to the final prediction. The output dimension of a MLP depends on the prediction task. For drug response prediction, the number of output dimensions is equal to the number of drugs, and for cancer type and subtype classification, the number of output dimensions is equal to the number of cancer types and subtypes.

### Model training

The sample size available for deep learning is limited, which poses a challenge for conducting a grid search. This approach typically involves reserving a portion of the dataset for final performance evaluation, further reducing the size of the training set. As a result, it will be difficult to achieve optimal model performance due to insufficient training data. For this reason, all methods were trained with default hyperparameters instead of hyperparameter tuning.

Default hyperparameters were used for random forest, elastic net, PCA (top 200 PCs) and *k*-NN (*k* = 5) and details can be found in the official API of scikit-learn (v1.0.2). Default hyperparameters were also set for mixOmics and moCluster, and details can be found in their original publications. The default hyperparameters of DeePathNet can be found in the GitHub repository. To train DeePathNet for regression, mean squared error (MSE) loss was computed between the predicted and actual IC_50_. For classification, we computed the cross-entropy (CE) loss to train DeePathNet.

The computation time of DeePathNet was reported in **Table S9** and was only compared with random forest and elastic net.

### Evaluation metrics

For regression, R^2^, MAE and Pearson’s *r* were used to evaluate the performance and they are defined as follows:

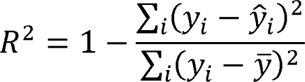

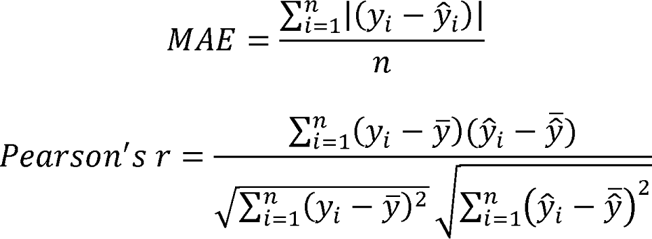

For a given drug, *y_i_* represents the actual IC_50_ of cell line *i*, *ŷ_i_* represents the predicted IC_50_ value of cell line *i*, *ȳ* represents the mean value of all actual IC_50_ values, *ŷ̄* represents the mean value of all predicted IC_50_ values, and *n* represents the total number of cell lines. For classification, multiple metrics were used to evaluate the predictive performance of DeePathNet and other models, including accuracy, macro-average F_1_-score, precision, recall, AUROC, AUPRC and stability. Let TP, TN, FP, FN represent true positive, true negative, false positive and false negative predictions. Accuracy is defined as 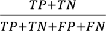. Precision is defined as 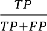. Recall is defined as 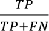. Then the F_1_-score is calculated as the harmonic mean of the precision and recall, defined as 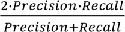. The macro-average F_1_-score is calculated by computing the arithmetic mean of F_1_-scores from all the cancer types or subtypes. The ROC curve is created by plotting the recall and false positive rate 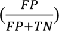 at various thresholds. AUROC is calculated as the area under the ROC curve. The precision and recall (PR) curve is created by plotting the precision and recall at various thresholds, and the AUPRC is calculated as the area under the PR curve. The stability is measured by the standard deviation.

## Results

### DeePathNet predicts drug response

We first assessed the predictive performance of DeePathNet on a regression task by benchmarking it against random forest [46], elastic net [47], principal component analysis (PCA), mixOmics [4], and moCluster [8] to predict the responses of anti-cancer drugs to cancer cell lines. These six methods were evaluated using data from the Cell Lines Project (CLP) [38] and the Cancer Cell Line Encyclopedia (CCLE) [39], the two largest publicly available multi-omic cancer cell line datasets (Table S1, see Experimental Procedures). DCell [20], DrugCell [21] and EMOGI [22] were not included in the benchmark as they do not support multi-omic data and general prediction tasks. Gene mutation, copy number variation (CNV), and gene expression data from the two datasets were used as the input. For drug response data, we retrieved the half-maximal inhibitory concentration (IC_50_) from the Genomics of Drug Sensitivity in Cancer (GDSC) [38] database. For each method, six experimental setups were assessed, comprising two datasets and three evaluation metrics, namely coefficient of determination (R^2^), mean absolute error (MAE), and Pearson correlation coefficient (Pearson’s *r*) between predicted and actual IC_50_ values.

In DeePathNet, 241 pathway encoders were constructed (Figure S1A) to summarize the omic data into pathway vectors defined by the LCPathways [26]. These vectors were then fed into the Transformer module to model the interactions among cancer pathways (Figure S1B). Default hyperparameters were used for all six methods (see Experimental Procedures). Omic data were combined using early integration [7] for random forest and elastic net. Middle integration [7] was used for PCA, moCluster, and mixOmics. PCA and moCluster were coupled with random forest for predictions [7] (see Experimental Procedures).

To quantitatively and reliably compare the six methods, five-fold cross-validation was repeated five times at random, yielding 25 error measures for each of the R^2^, MAE, and Pearson’s *r* metrics. The mean and 95% confidence interval (CI) of the evaluation metrics was reported, serving as an estimate of the generalization error. We observed that DeePathNet had significant and consistently better performance in drug response prediction than the other five methods that do not incorporate cancer pathway information (Figure 2A-F, *p*-value < 0.0001, two-tail paired Student’s *t*-test, Table S2). By ranking the methods according to the mean measures for each setup, we found that random forest was the second-best performing method (Figure 2G). To investigate whether drug responses that had relatively lower predictive accuracy by DeePathNet were also challenging for other methods, the correlations between DeePathNet and the other five methods were calculated. We found that the predictive performance of the paired methods was highly concordant (Pearson’s *r* > 0.9), with DeePathNet consistently outperforming the other five methods (Figure S2).

**Figure 2.**
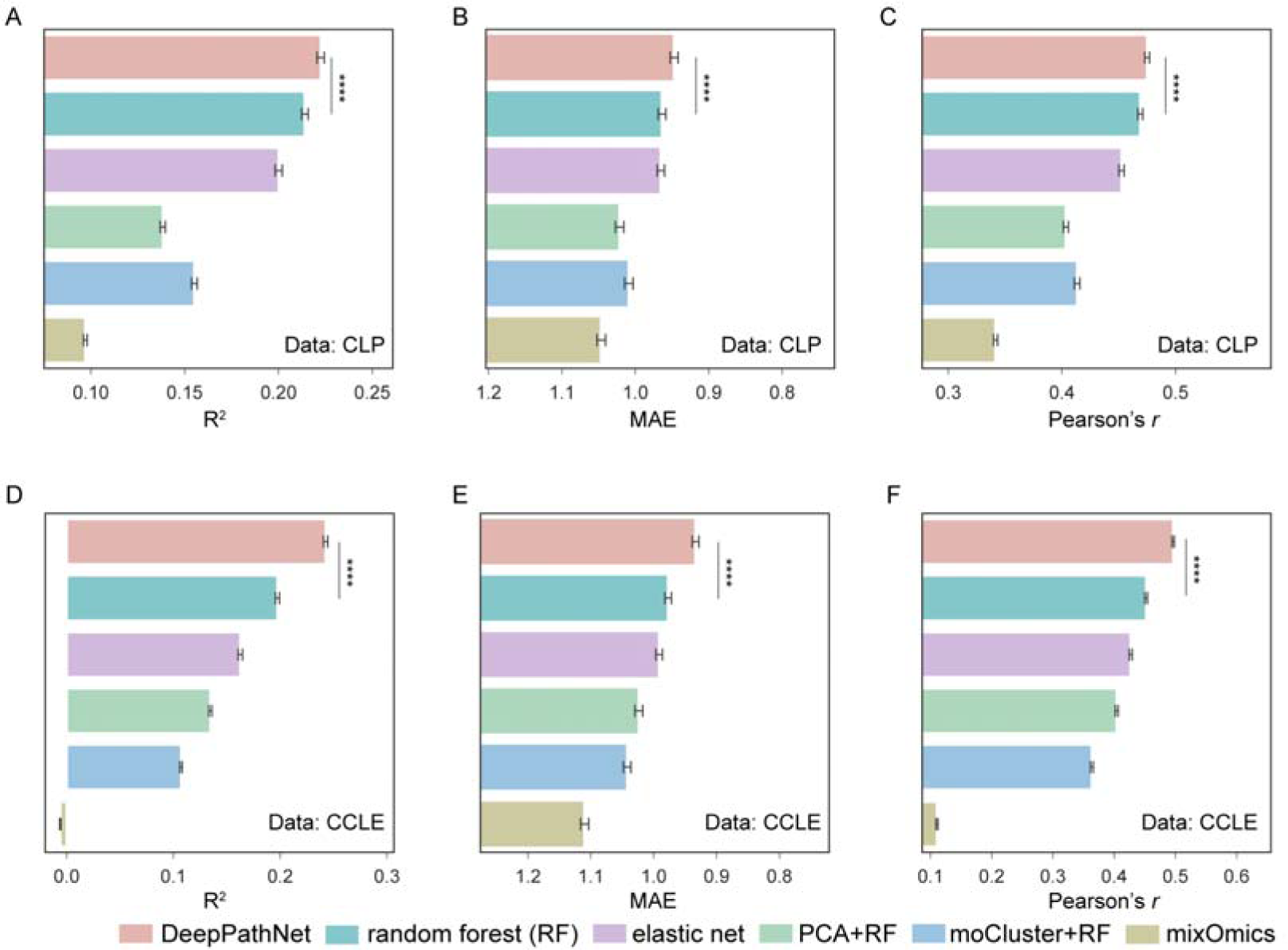
Performance evaluation of drug response prediction by cross-validation. **A-F,** Bar plots showing predictive performances across six experimental setups on the CLP and CCLE datasets by three evaluation metrics: R^2^, MAE (inverted on the horizontal axis), and Pearson’s *r*. A higher value represents better performance. Error bars are derived from cross-validation, representing 95% confidence intervals of the mean. **** indicates *p*-value < 0.0001 by two-tail paired Student’s *t*-test, only showing significance between the first-and second-best performing methods. The six methods are color-coded at the bottom.

To validate the model’s performance using an independent test set, we trained DeePathNet on the CLP dataset (Table S1) and tested the final model by predicting drug responses in the CCLE dataset (Table S1). Overlapping cell lines between CLP and CCLE were excluded from the test set (see Experimental Procedures). Cancer pathway information was integrated in the same way as described above, and a random forest model was trained as the baseline. The test performance for all 549 GDSC anti-cancer drugs was summarized for both DeePathNet and random forest. DeePathNet achieved statistically significantly higher predictive performance than random forest across all three metrics (Figure 3A, *p*-value < 0.0001, two-tail paired Student’s *t*-test, Table S3).

**Figure 3.**
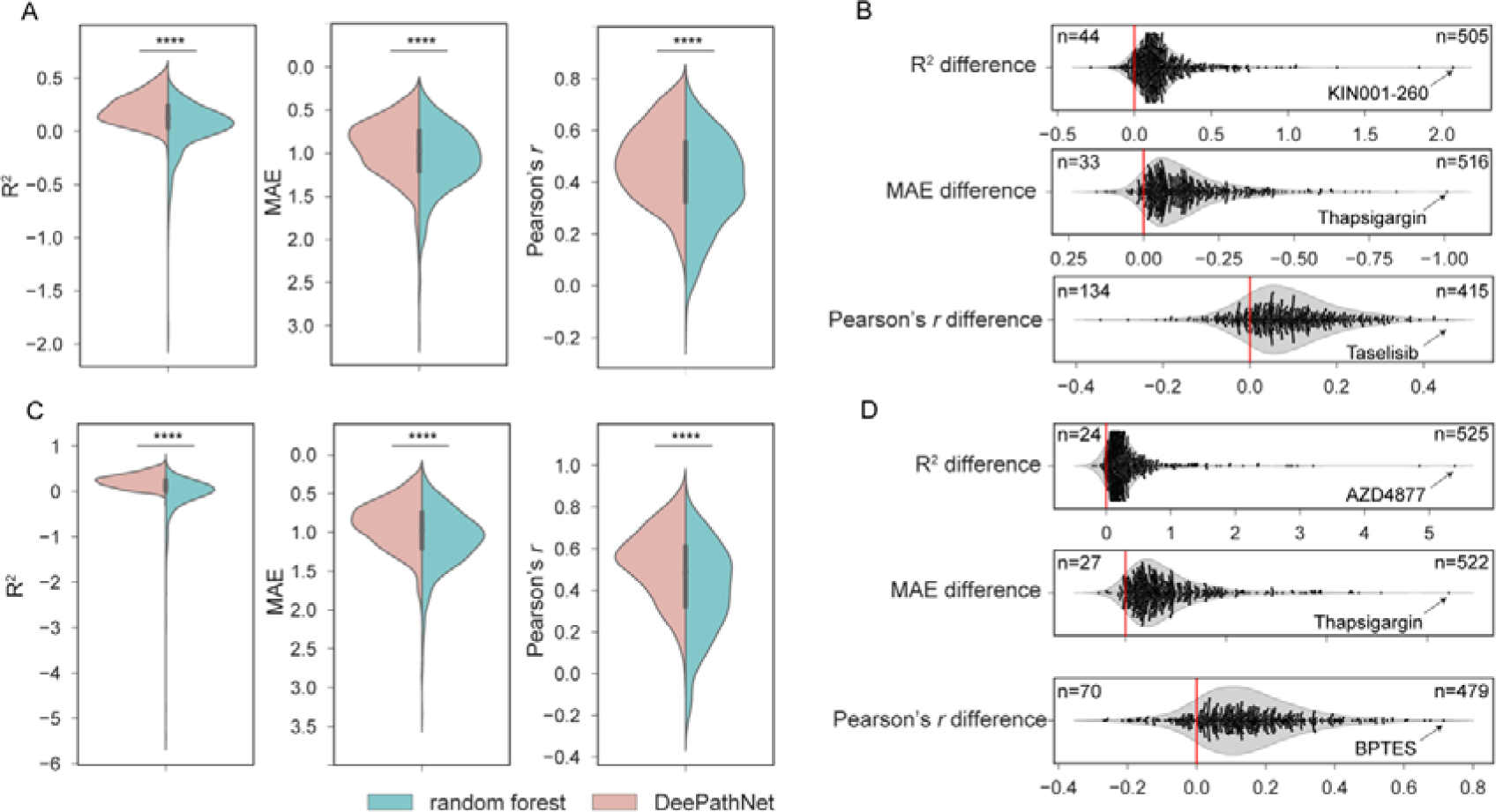
Generalization error of DeePathNet and random forest for drug response prediction. **A,** Violin plots showing predictive performances of DeePathNet and random forest using CLP as the training set and evaluated on the independent CCLE test set across the 549 GDSC drugs. The vertical axis is inverted for MAE. **** indicates *p*-value < 0.0001 by two-tail paired Student’s *t*-test. **B,** Violin and swarm plots showing the performance difference in R^2^ (upper), MAE (middle) and Pearson’s *r* (lower) between DeePathNet and random forest for each drug. A drug is more accurately predicted by DeePathNet when it exhibits a positive value for the R^2^ or Pearson’s *r* difference, or a negative value of the MAE difference (the horizontal axis is inverted for MAE). The numbers of drugs more accurately predicted by DeePathNet or random forest are annotated on the upper right and left of the plot, respectively. The name of the drug that achieved the largest improvement with DeePathNet is annotated for each metric. **C** and **D**, Similar to **A** and **B**, but using CLP^+^ as the training set and CCLE^+^ as the independent test set.

To compare the predictive performance of DeePathNet with random forest for each drug, the difference of R^2^ between DeePathNet and random forest was measured. Here, 92% (505/549) of drugs had positive values, indicating superior predictive performance from DeePathNet over random forest. Similarly, 94% (516/549) of drugs and 76% (415/549) of drugs exhibited improved results by MAE and Pearson’s *r*, respectively (Figure 3B). This demonstrates that DeePathNet consistently achieved better predictive performance than random forest for most anti-cancer drugs. The drug that obtained the largest R^2^ improvement by DeePathNet was KIN001-260 (Figure 3B), which was poorly predicted by random forest and caused a long tail in the distribution of R^2^ values (Figure 3A). Drugs that had the largest improvement in MAE and Pearson’s *r* with DeePathNet were thapsigargin and taselisib (Figure 3B).

Next, we extended our analysis by including two proteomic cell line datasets from ProCan-DepMapSanger [40] and CCLE [41]. ProCan-DepMapSanger is a recently published pan-cancer proteomic dataset of 949 human cell lines generated by our team, supplementing the CLP with proteomic information. DeePathNet and random forest were trained on the combined CLP and ProCan-DepMapSanger datasets (CLP^+^; Table S1), with the final model tested on the expanded CCLE dataset that includes additional proteomic measurements (CCLE^+^; Table S1). Pathway information was integrated into DeePathNet as described above. DeePathNet yielded significantly higher test performance than random forest across all three metrics when predicting the 549 GDSC anti-cancer drugs (Figure 3C, Table S3). Analyzing the predictive performance for each drug, DeePathNet also provided significant improvement for the majority of anti-cancer drugs compared with random forest (Figure 3D). The drugs that had the largest improvement by DeePathNet were AZD4877, thapsigargin and BPTES measured by the differences of R^2^, MAE and Pearson’s *r*, respectively (Figure 3D).

To investigate which types of drugs were most accurately predicted by DeePathNet, we grouped the 549 drugs by their canonical target cellular pathways. Drugs targeting ABL signaling and ERK MAPK signaling pathway had the highest mean Pearson’s *r* between predicted and actual IC_50_ values (Figure S3A). The top 20 most accurately predicted drugs and their pathways are reported in Figure S3B.

Taking these observations together, we demonstrated that DeePathNet increased predictive performance through several benchmarking analyses to predict responses to several drugs targeting various signaling pathways. The performance of DeePathNet on drug response prediction was also validated using independent datasets.

### DeePathNet classifies cancer types

To evaluate DeePathNet on a classification task, we used publicly available data from The Cancer Genome Atlas (TCGA) [48] to classify primary cancer types. Gene mutation, CNV and gene expression features were used as the omic data input to train DeePathNet models to classify each of the 6,356 samples into one of 23 cancer types (see Experimental Procedures). A total of seven metrics were used across the analyses to ensure reliable evaluation. These metrics are accuracy, macro-average F_1_-score, precision, recall (sensitivity), area under the receiver operating characteristic curve (AUROC), area under the precision-recall curve (AUPRC) and stability (see Experimental Procedures). LCPathways was integrated in the same way as described for the drug response prediction. For benchmarking, the elastic net was replaced by the *k*-nearest neighbors (*k*-NN) [49]. This was done because elastic net regularization is less commonly used for the task of cancer type classification, while *k*-NN is a more widely adopted method for classification. For all six methods, feature integration and hyperparameter settings were identical to the drug response prediction.

In the absence of an independent dataset comprising the 23 cancer types, cross-validation was performed for these six methods on the TCGA dataset, and the mean and 95% CI of the evaluation metrics were reported as an estimate of the generalization error. DeePathNet consistently outperformed the other five machine learning methods by accuracy and macro-average F_1_-score (Figure 4A, Table S4). In contrast, other methods, such as mixOmics, only performed well in one metric, indicating that these methods may be suitable for certain scenarios but cannot generalize well across different prediction tasks (Figure 4A). Assessing the performance of each method using a set of four metrics including accuracy, macro-average F_1_-score, AUROC and stability showed that DeePathNet was consistently top-ranked, followed by random forest (Figure 4B, Table S4).

**Figure 4.**
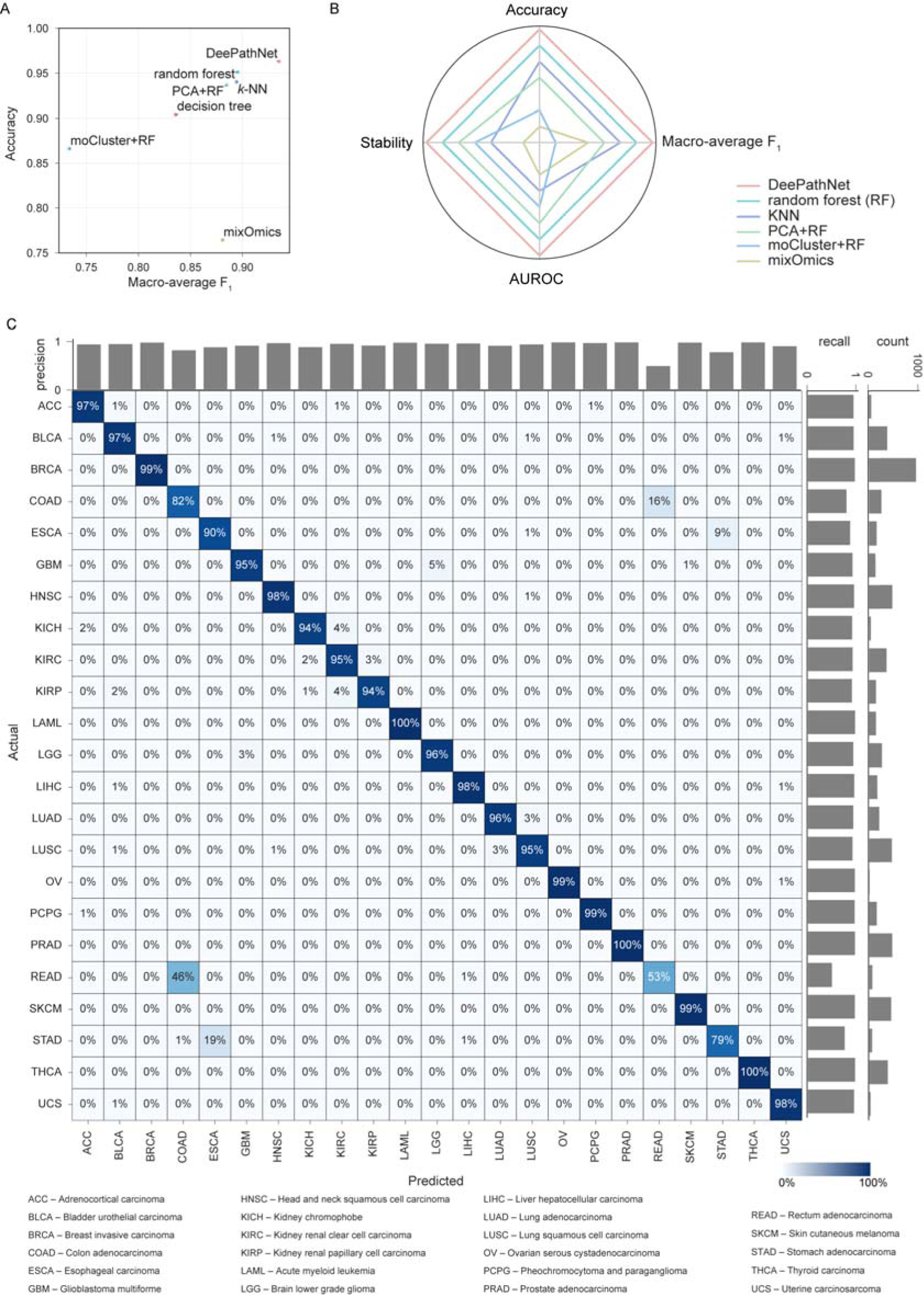
Performance evaluation of cancer type classification. **A,** Model comparison using cross-validation on the TCGA dataset. The x-axis represents the macro-average F_1_-score, and the y-axis denotes accuracy. **B,** Radar chart showing the model ranks across the set of four metrics. A larger enclosed area indicates better predictive performance. **C,** Confusion matrix for the classification of 23 cancer types. Columns denote predicted labels, and rows represent actual labels. The percentage shown represents the proportion of predictions made for the corresponding cancer type, with each row summing to 1. The diagonal represents correct predictions for each cancer type, with the percentage indicating the recall. Bar plots show precision (horizontal axis), recall (vertical axis, leftmost) and number of samples (vertical axis, rightmost) per cancer type.

To further investigate DeePathNet’s performance for each cancer type, the predicted and actual cancer type for each sample was visualized using a confusion matrix, with the number of samples, precision and recall annotated (Figure 4C). DeePathNet achieved a recall of over 0.95 for most cancer types, with acute myeloid leukemia (LAML), pancreatic adenocarcinoma (PRAD), and thyroid carcinoma (THCA) as the top three most accurately classified cancer types. Rectum adenocarcinoma (READ) was the cancer type with the lowest recall, with 46% of the samples incorrectly classified as colon adenocarcinoma (COAD). The latter outcome is unsurprising because the colon and rectum are adjacent tissue types that share highly similar features, with these two cancer types often grouped together [50] and are treated with similar chemotherapeutic regimens [50]. The cancer type exhibiting the second-lowest recall was stomach adenocarcinomas (STAD), with 19% of STAD samples incorrectly classified as esophageal carcinoma (ESCA). This can be explained by their similar histopathology and the anatomical proximity of STAD and ESCA [51]. Next, AUROC and

AUPRC were examined for each cancer type, both displaying high performances for all cancer types, with the exception of AUPRC for READ, due to the tissue proximity of READ to COAD. (Figure S4A and Figure S4B).

### DeePathNet classifies breast cancer subtypes

Gene mutation, CNV and gene expression features were used to train DeePathNet models for the classification of five breast cancer subtypes (Luminal A, Luminal B, HER2+, Basal, Normal-like) according to the Prediction Analysis of Microarray 50 (PAM50) test [52]. A total of 974 breast cancer samples from the TCGA dataset were used for training, and a breast cancer cohort of 122 samples from the Clinical Proteomic Tumor Analysis Consortium (CPTAC) was included as an independent dataset to evaluate the generalization error.

Cross-validation for all six methods was first performed on the TCGA dataset, reporting the mean and 95% CI of the evaluation metrics as an estimate of the generalization error. DeePathNet substantially improved over the other methods in terms of accuracy and macro average F_1_-score (Figure 5A, Table S5). The performance gain in AUROC was relatively minor but statistically significant (Student’s *t*-test *p*-value < 5x 10^-4^) (Table S5). The methods were then ranked according to the same set of four metrics as in cancer type classification. DeePathNet achieved the best performance in all four metrics, with random forest ranked as the second best overall (Figure 5B). Other methods showed inconsistent performance rankings across different metrics, demonstrating the necessity of using multiple evaluation metrics for a comprehensive evaluation.

**Figure 5.**
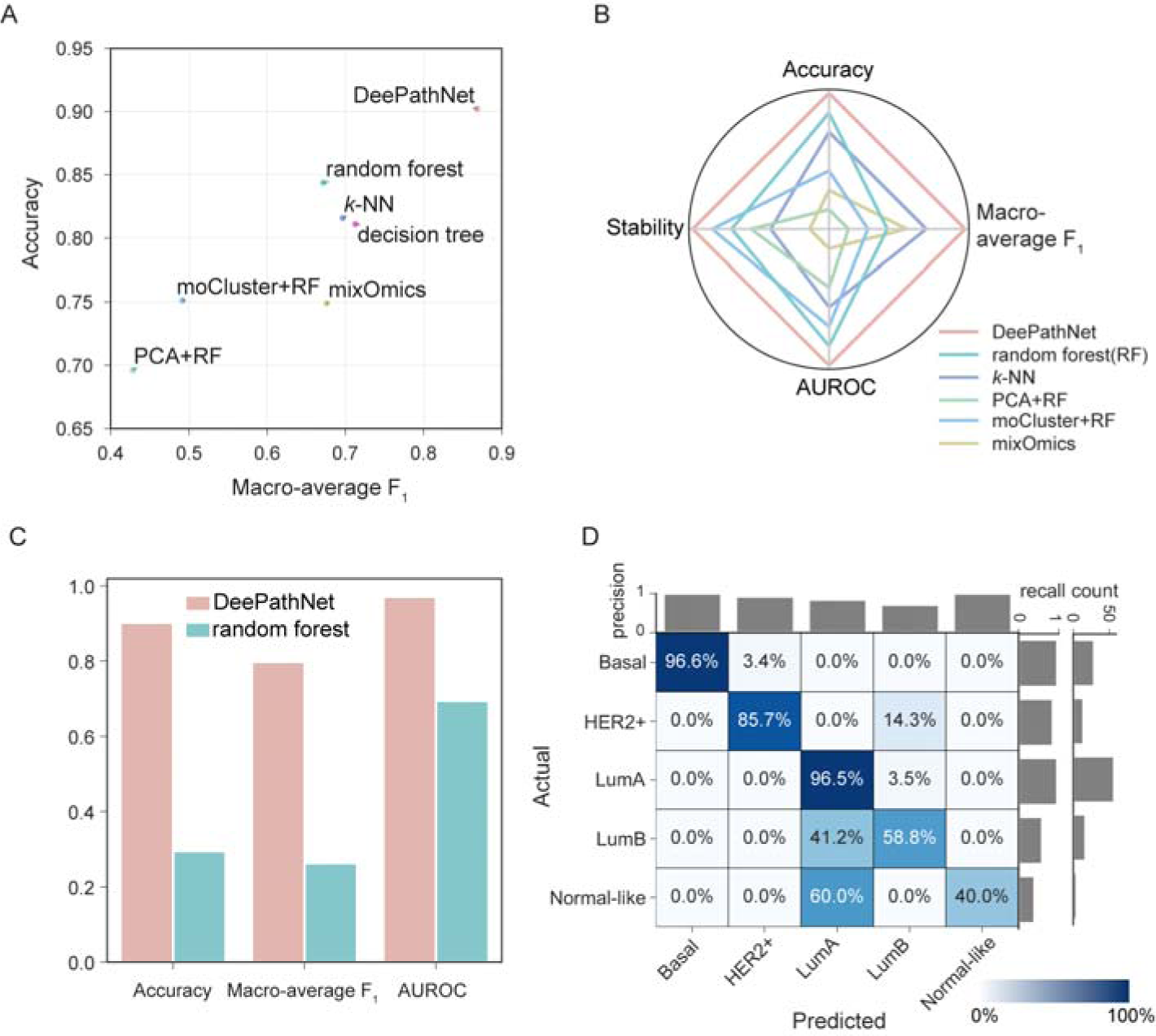
Performance evaluation of breast cancer subtype classification. **A,** Model evaluation by cross-validation. The x-axis represents the macro-average F_1_-score, and the y-axis represents accuracy. **B,** Radar chart showing the model ranks across the set of four metrics. A larger enclosed area represents better classification performance. **C,** Performance metrics showing generalization errors for DeePathNet and random forest when using CPTAC data as the independent test set for validation. **D,** Confusion matrix showing generalization errors when using CPTAC data as the independent test set for validation. Statistics are annotated in the same way as described in Figure 4C.

To validate the model’s performance on an independent test set, a DeePathNet model was trained on the TCGA breast cancer cohort, with the final model tested on the independent CPTAC breast cancer cohort. Benchmarked against random forest, DeePathNet yielded a much lower generalization error on the independent test set (Figure 5C, Table S6). Next, the generalization error of DeePathNet was assessed for each subtype by a confusion matrix. DeePathNet achieved the highest precision and recall in classifying the Basal subtype (96.6%, Figure 5D), with most tumors in this subtype being high-grade with a poor prognosis [53]. The most difficult subtype to classify was Normal-like, where three out of the five Normal-like samples were incorrectly classified as Luminal A (Figure 5D). Luminal A and Normal-like subtypes are traditionally difficult to distinguish as they share the same immunohistochemistry markers [53]. The Normal-like subtype is less frequently used in clinics [54]. Further analyses by AUROC (Figure S5A) and AUPRC (Figure S5B) demonstrated DeePathNet’s high predictive performance for each subtype. Overall, these results demonstrated the performance of DeePathNet on cancer subtype classification using both cross-validation and validation using an independent dataset.

### DeePathNet provides model explanations

The DeePathNet model is explainable at both omic and pathway levels by using feature importance derived from SHapley Additive exPlanations (SHAP) [55] and Layer-wise Relevance Propagation (LRP) [56]. SHAP attributes the prediction to all features, assigning each feature an importance value. At the same time, LRP assumes that the classifier can be decomposed into several layers of computation, with these layers being parts of the feature extraction. Thus, SHAP and LRP are post-hoc model explanation approaches that establish relationships between feature values and the predictions after DeePathNet is trained. Breast cancer subtype classification was used to demonstrate the model explanation.

To explain the model at the omic level, SHAP was used to calculate feature importance. Specifically, feature importance was computed and visualized for the top five genes as stack bar plots comprising each omic data type for each breast subtype (Figure 6A). DeePathNet was able to identify known biomarker genes as top features, such as ESR1, ERBB2 and KRT17, whose gene expression is routinely used to determine the PAM50 subtypes in the clinic (Figure 6A) [52]. Most genes had their high feature importance attributable to transcriptomic data (Figure 6A), consistent with the fact that PAM50 classifications are RNA-based subtypes [52].

**Figure 6.**
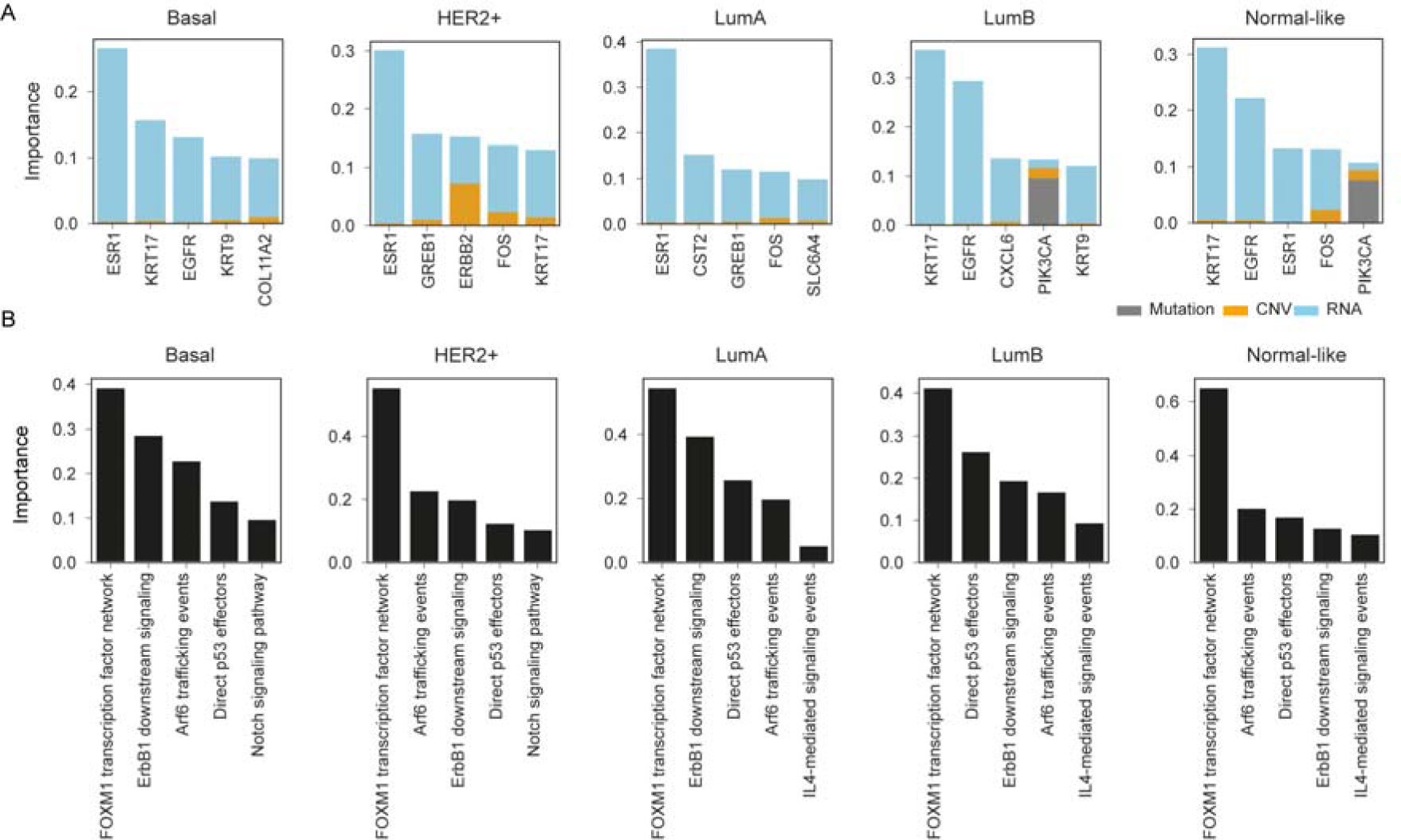
DeePathNet model explanation by omic level and pathway level feature importance. **A,** Stacked bar plots showing the omic level feature importance of the top five genes for each omic data type (indicated by grey, yellow, and blue color). **B,** Bar plots showing the DeePathNet pathway level feature importance of the top five pathways.

To explain the models at the pathway level, LRP was used to calculate feature importance. Since the cancer pathways are represented as an encoded vector that summarizes multi-omic information, the feature importance of a cancer pathway is computed for all omic data types jointly. For each cancer subtype, the top five pathways with the highest feature importance values were ranked (Figure 6B). DeePathNet identified the FOXM1 transcription factor network as the most important pathway for predicting all PAM50 subtypes (Figure 6B). FOXM1 shows distinct patterns of expression in different breast cancer subtypes and is seen as a promising candidate target in breast cancer treatment [57]. FOXM1 is also an adverse prognostic factor of survival in Luminal A and B subtypes [58]. The ARF6 pathway was shown to be overexpressed in triple-negative breast cancer and to be associated with breast cancer invasion and metastasis [59]. Similarly, Notch Signaling pathways are involved in cell proliferation, apoptosis, hypoxia and epithelial to mesenchymal transition and were found to be over-expressed in HER2 positive and triple-negative breast cancer [60].

Taken together, these findings suggest that DeePathNet provides reliable model explanations with a strong biological basis by providing feature importance at both the omic level and pathway level.

### Ablation study

Lastly, we assessed the predictive capability of DeePathNet by separately examining its two primary components: the pathway encoder and the Transformer module. To evaluate the additional predictive strength by integrating cancer pathway knowledge, we devised a control by randomly altering the pathway definitions for DeePathNet. This process was repeated tenfold. Compared with the random control, DeePathNet provided more accurate predictions across all the metrics for breast cancer subtype classification, while similar outcomes were observed for drug response prediction and cancer type classification (Figure 7A-C, Table S7 and S8). Such comparable performance between randomly configured and manually designed neural networks was also observed previously [21,61]. Without losing any predictive power, neural networks designed using human knowledge have been found to provide more meaningful model explanations that could lead to biological discoveries [21,61]. Further investigation replicated the phenomenon observed in the previous study [61] demonstrating that the neural network, constructed using known biological knowledge, outperforms random control when the sample size is limited (Figure 7D). Next, we benchmarked DeePathNet using a multilayer perceptron (MLP) as a plain neural network that does not use either the pathway information or the Transformer. When equipped with both the pathway information and the Transformer module, DeePathNet achieved significantly more accurate predictions across all considered metrics and tasks (Figure 7A-C, Table S7 and S8).

**Figure 7.**
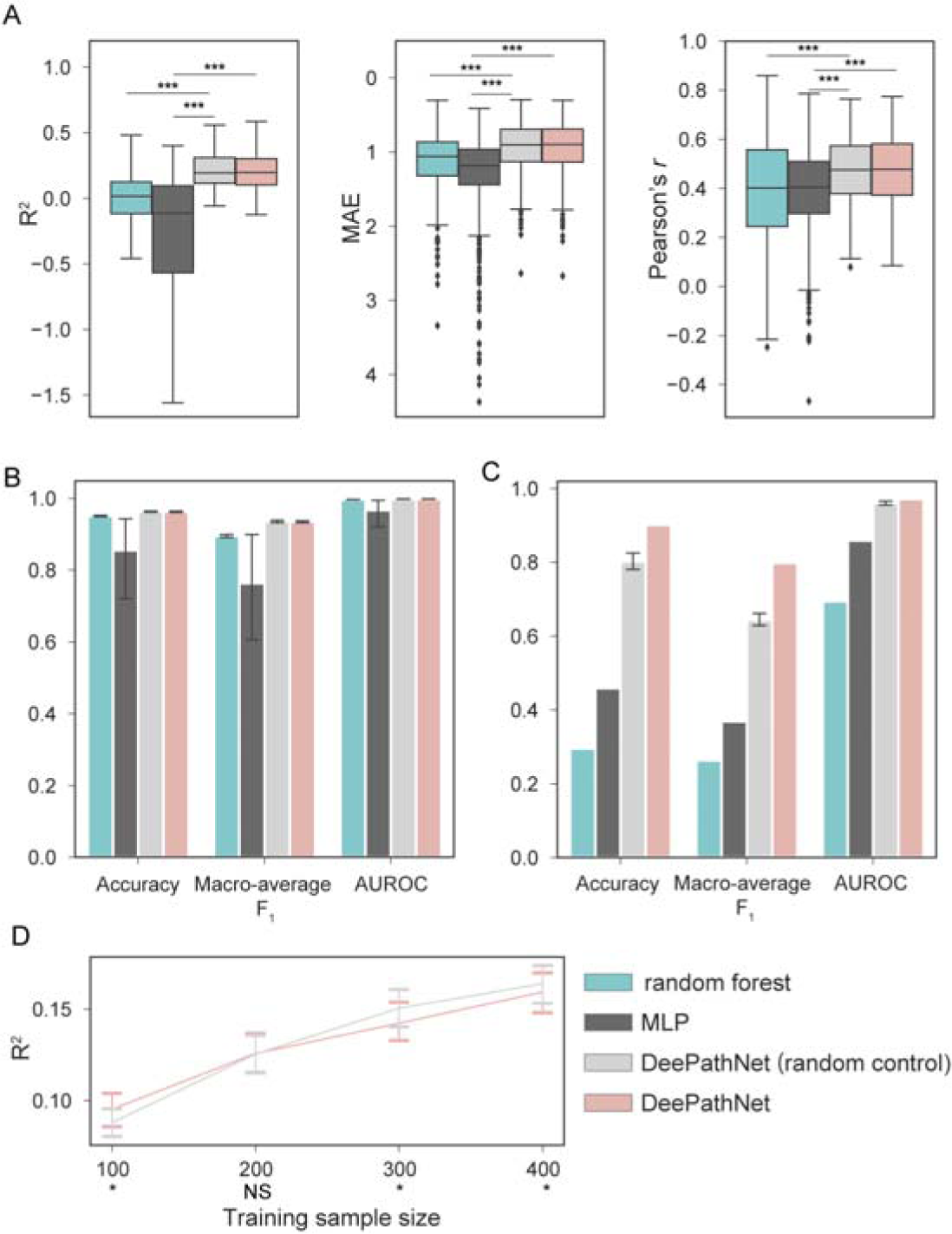
Ablation study of DeePathNet using randomly wired pathways (random control) and plain neural networks (MLP). **A,** Box plots showing the predictive performance from the independent test set across the 549 drugs for random forest (green), MLP (dark grey), DeePathNet with randomly wired pathways (light grey) and DeePathNet (pink). Outliers for R^2^ are hidden for clearer visualization. **B,** Bar plots showing predictive performances of the four models from cross-validation on cancer type classification. **C.** Bar plots showing predictive performance from the independent test set on breast cancer subtype classification. Confidence intervals represent 95% confidence intervals of the mean (n = 10 experiments). **D.** Down-sample analysis showing DeePathNet with cancer pathways performs better on drug response prediction with smaller training sample sizes compared with DeePathNet with random wired pathways. The solid line represents the mean R^2^, and confidence intervals represent 95% confidence intervals of the mean (n = 10 experiments) \**P* < 0.0001, paired *t*-test.

In summary, DeePathNet, when integrated with the Transformer module – irrespective of the presence of pathway knowledge – provided significantly higher predictive power compared with plain neural networks and random forest. The inclusion of pathway knowledge enhanced DeePathNet’s accuracy in breast cancer subtype classification and facilitated pathway level model explanation.

## Conclusion

DeePathNet broadens the application of the Transformer to cancer multi-omic data integration by incorporating cancer pathways, which successfully overcomes the limitations of most existing machine learning approaches that do not consider known cancer biology. DeePathNet integrates multi-omic data with cancer pathway knowledge to accurately predict drug responses and classify cancer types and subtypes. The self-attention mechanism of the Transformer module dynamically models the interdependency among pathways, thus capturing regulatory effects across different biological processes and the effects of dysregulation. By modeling pathways instead of multi-omic features directly, DeePathNet also alleviates the challenge of the curse of dimensionality [44] commonly encountered in multi-omic data analysis.

The predictive performance of DeePathNet was evaluated and validated by one regression and two classification tasks. The evaluation was conducted on a larger scale than previous similar studies [8,11] using multiple big datasets and a range of metrics with both cross-validation and independent testing. Based on the Transformer architecture, DeePathNet outperformed other machine learning methods that do not incorporate pathway information and instead rely only on omic data for input features. A low generalization error when validating DeePathNet models on independent datasets suggests that DeePathNet works well even when different experimental protocols were implemented among these independent datasets. DeePathNet provides model explanations at the pathway level, which has not yet been accomplished by any other multi-omic integration tools that can predict drug response or classify cancer type and subtype. DeePathNet was able to highlight known biomarkers when predicting breast cancer subtypes, including ESR1, ERBB2 and the FOXM1 network pathways. This suggests that other top-ranked genes and pathways may provide novel insights into cancer biology and drug discovery.

Despite these comprehensive evaluations, DeePathNet has only been designed to support gene-based features at this stage. Other omic data types such as phosphoproteomics, metabolomics and images cannot be directly utilized as the input to the pathway encoder. Therefore, future work of DeePathNet includes extending the model to support any data modalities, which can be potentially done by mapping the features into the pathway space using different types of encoders. Meanwhile, as large proteomic and metabolomic datasets become increasingly available, the predictive power of DeePathNet will improve because deep learning will obtain a performance boost with more data [62].

In conclusion, DeePathNet combines multi-omics, deep learning, and existing biological knowledge to predict cancer phenotypes accurately with a model explanation. The application of DeePathNet may lead to more accurate diagnosis and prognosis and will facilitate researchers to understand unknown cancer mechanisms and prioritize putative drug targets.

## Acknowledgements

ProCan® is supported by the Australian Cancer Research Foundation, Cancer Institute New South Wales (NSW) (2017/TPG001,REG171150), NSW Ministry of Health (CMP-01), The University of Sydney, Cancer Council NSW (IG 18-01), Ian Potter Foundation, the Medical Research Futures Fund (MRFF-PD), National Health and Medical Research Council (NHMRC) of Australia European Union grant (GNT1170739, a companion grant to support the European Commission’s Horizon 2020 Program, H2020-SC1-DTH-2018-1, ‘iPC-individualized Paediatric Cure’ [ref. 826121]), and National Breast Cancer Foundation (IIRS-18-164). Work at ProCan® is done under the auspices of a Memorandum of Understanding between Children’s Medical Research Institute and the U.S. National Cancer Institute’s International Cancer Proteogenome Consortium (ICPC), that encourages cooperation among institutions and nations in proteogenomic cancer research in which datasets are made available to the public. Z.C. is the recipient of a PhD Scholarship from Sydney Cancer Partners with funding from Cancer Institute NSW (2021/CBG0002). R.C.P. is supported by the NHMRC of Australia (GNT1138536 and GNT2000855). Some figures were created with BioRender.com.

## Author Contributions

Z.C. and Q.Z. conceived and designed the experiments and wrote the manuscript. Z.C. analyzed the data and developed the method. R.C.P., A.A., P.J.R. and R.R.R. contributed to the manuscript.

## Conflicts of Interest

The authors declare no conflict of interest.

## Data and Code Availability

All data used in this study are publicly available datasets. Data can be downloaded from the original publication as cited. The source code and documentation of DeePathNet is available at https://github.com/CMRI-ProCan/DeePathNet. Source code and intermediate files are also available at https://doi.org/10.6084/m9.figshare.24137619

## Notes

### Competing Interest Statement

The authors have declared no competing interest.

### Summary of Updates

updated a few figures and texts

