## Supplementary Information for "Transformer-based deep learning integrates multi-omic data with cancer pathways"

**Notes:**

* Correspondence.

**Keywords:**

Multi-omics; Deep Learning; Cancer; Model Explanation, Cancer Pathway

### Supplementary Figures

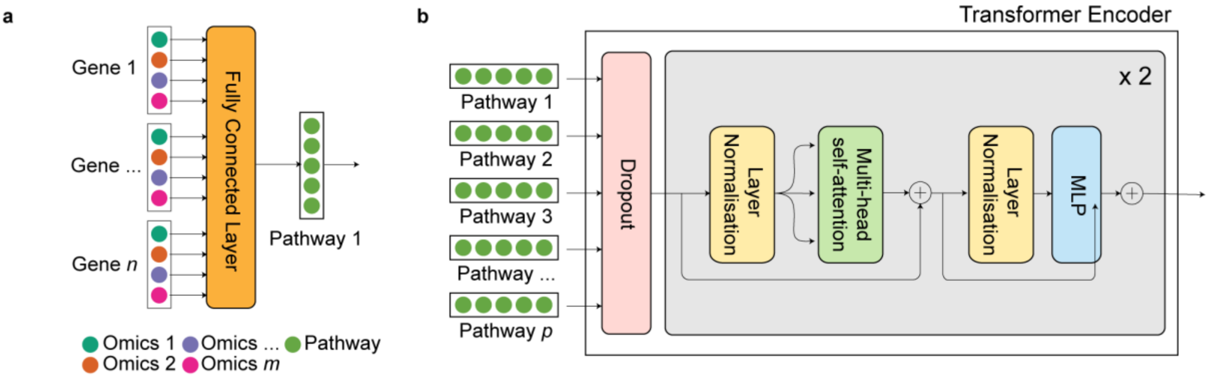

**Supplementary Figure 1 | Details of pathway encoder and Transformer encoder. a,** Detailed illustration of the pathway encoder. A fully connected layer encodes multi-omic data from genes into a pathway vector. **b,** Detailed illustration of the Transformer encoder. Pathway vectors are first fed into a dropout layer, followed by a recurrent sequence (grey box) of layer normalisation, multi-head self-attention and multi-layer perceptron (MLP). The components in the grey box recur twice in DeePathNet. Arrows represent the direction of information flow and ⊕ represents matrix addition.

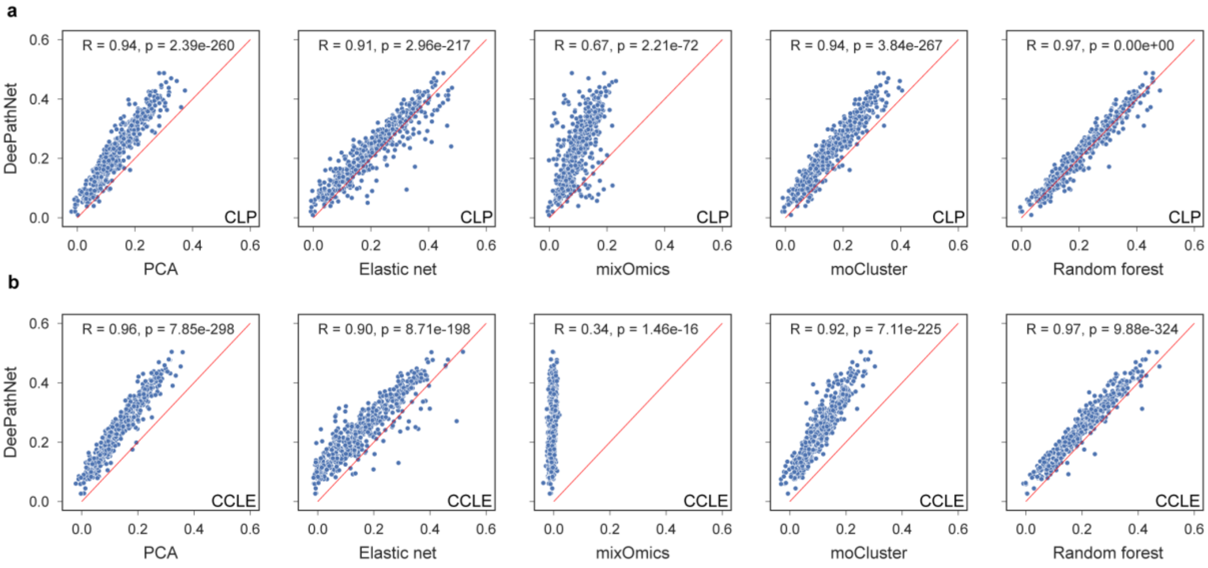

**Supplementary Figure 2 | Consistency of drug response predictions from different machine learning models. a,** Scatter plots showing $R^2$ of drugs from DeePathNet (vertical axis) against each of the remaining five machine learning models (horizontal axis) evaluated on the CLP dataset. The diagonal red line indicates equal performance between the two models. Points above the red line represent drugs that are more accurately predicted by DeePathNet. Pearson's *r* (R) and *p*-value (p) are annotated. **b,** Similar to **a**, but evaluated on the CCLE dataset.

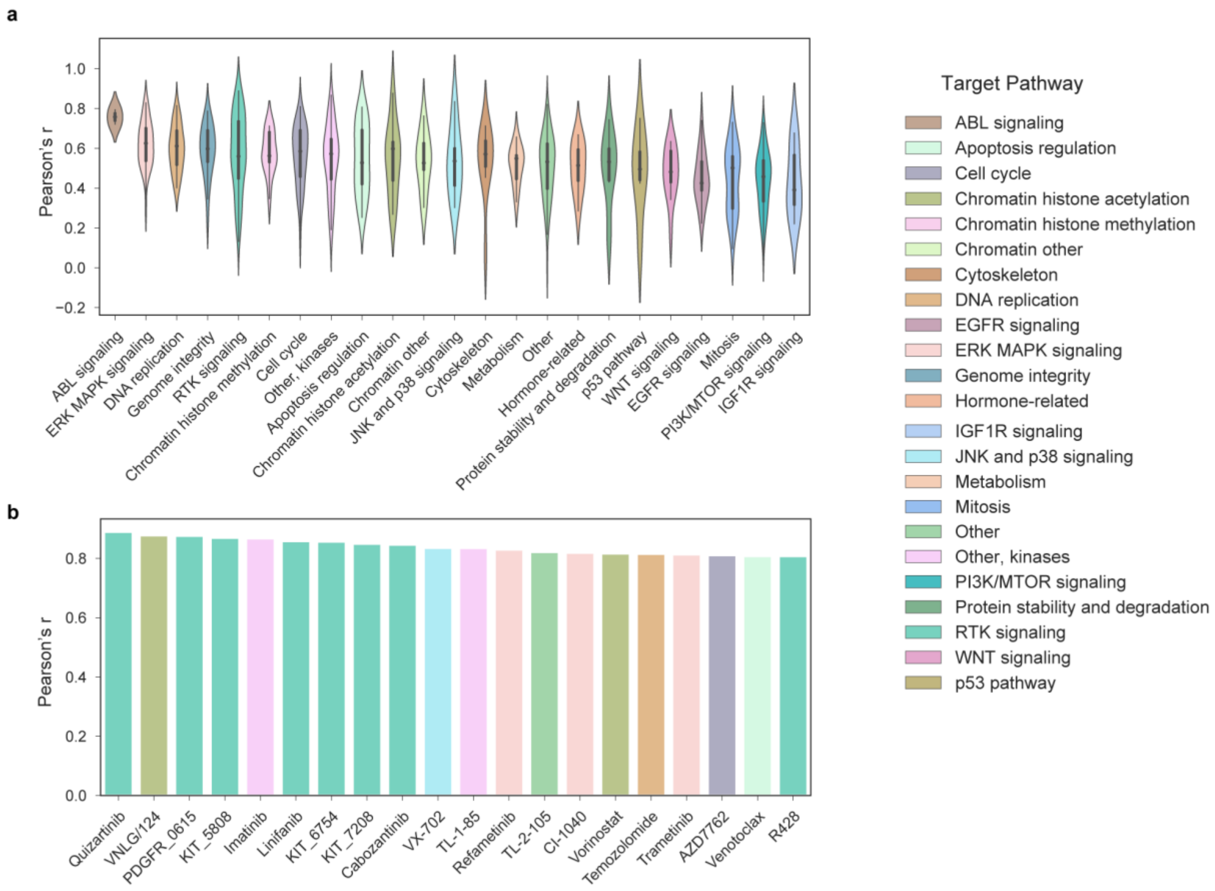

**Supplementary Figure 3 | Analysis of performance of drug response prediction by target pathways.** **a,** A violin plot showing the predictive performance grouped by drug canonical target pathways, ranked by the mean Pearson's *r* of the group. **b,** Top 20 drugs ranked by Pearson's *r*. Drugs are coloured by their canonical target pathways.

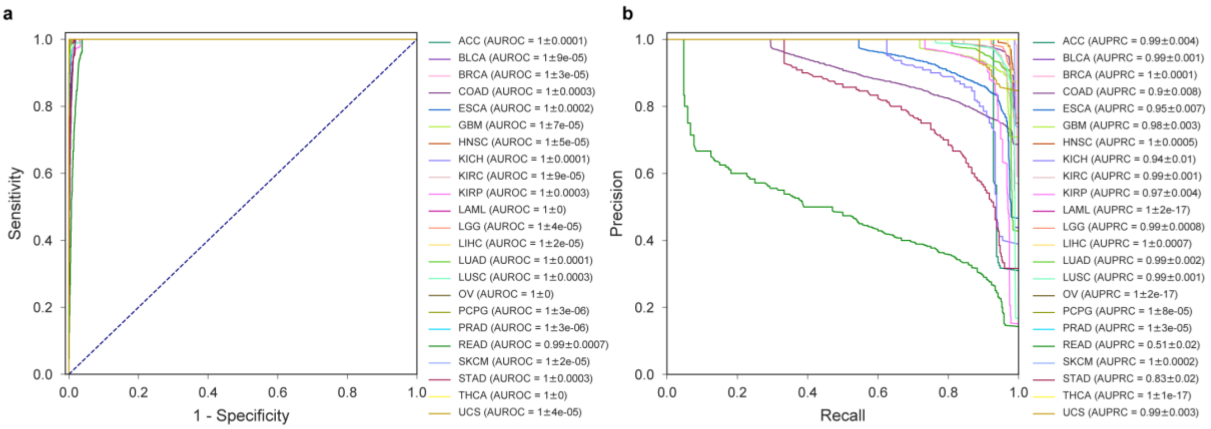

**Supplementary Figure 4 | ROC curves and precision-recall curves for TCGA cancer type classification. a,** ROC curves for DeePathNet classification of TCGA cancer types. Mean AUROC and standard error of the mean are annotated. **b,** Precision-recall curves for DeePathNet classification of TCGA cancer types. Mean AUPRC and standard error of the mean are annotated. Full terms of the abbreviations in **a** and **b** are listed in **Fig. 4c**.

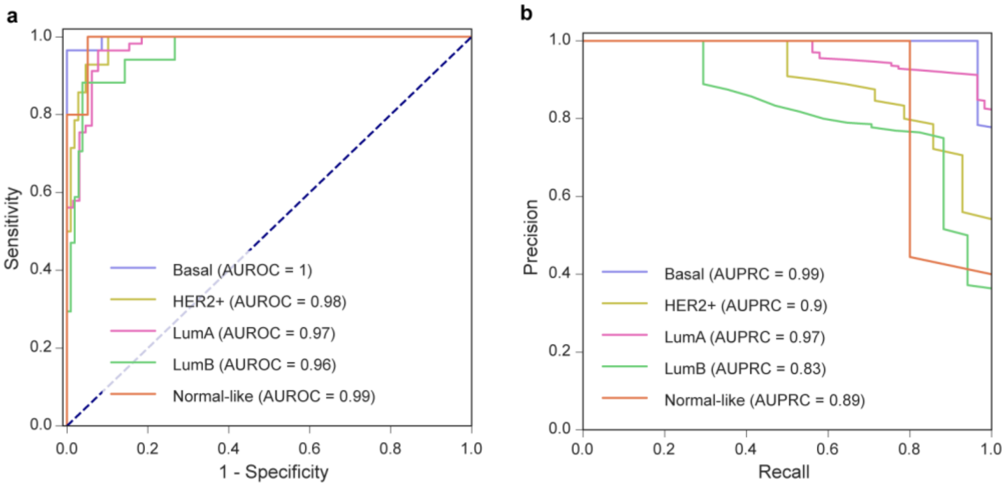

**Supplementary Figure 5 | ROC curves and precision-recall curves for breast cancer subtype classification. a,** ROC curves for DeePathNet classification of breast cancer subtypes using TCGA as the training data and CPTAC as the test data. **b,** Similar to **a**, but showing AUPRC.

### Supplementary Tables

| Data sets | Number of features | | | |
|---|---|---|---|---|
| (n = sample size) | **Gene mutation** | **CNV** | **Gene expression** | **Protein** |
| **Drug response prediction (n = 549 GDSC drugs)** | | | | |
| CLP (n = 941) | 19,099 | 19,116 | 15,320 | - |
| $CLP^+$ (n = 910) | 19,099 | 19,116 | 15,320 | 8,498 |
| CCLE (n = 696) | 18,103 | 27,562 | 19,117 | - |
| $CCLE^+$ (n = 292) | 18,103 | 27,562 | 19,117 | 12,755 |
| **Cancer type classification** | | | | |
| TCGA (n = 6,356) | 31,949 | 23,529 | 20,435 | - |
| **Breast cancer subtype classification** | | | | |
| TCGA (n = 974) | 31,949 | 23,529 | 20,435 | - |
| CPTAC (n = 122) | 11,877 | 23,692 | 23,121 | - |

**Supplementary Table 1.** Overview of datasets used in this study. $CLP^+$ = CLP + ProCan-DepMapSanger and $CCLE^+$= CCLE + CCLE proteomic dataset.

| | R2 mean ± 95%CI | MAE mean ± 95%CI | Pearson's r mean ± 95%CI |
|---|---|---|---|
| **CLP** | | | |
| **DeePathNet** | **0.222±0.0020** | **0.947±0.0021** | **0.475±00.0053** |
| Random forest (RF) | 0.214±0.0018 | 0.964±0.0021 | 0.469±0.0054 |
| Elastic net | 0.200±0.0020 | 0.965±0.0023 | 0.452±0.0051 |
| moCluster+RF | 0.155±0.0015 | 1.009±0.0022 | 0.413±0.0057 |
| PCA+RF | 0.138±0.0014 | 1.021±0.0022 | 0.403±0.0058 |
| mixOmics | 0.097±0.0009 | 1.047±0.0019 | 0.342±0.0061 |
| **CCLE** | | | |
| **DeePathNet** | **0.242±0.0020** | **0.934±0.0020** | **0.496±0.0052** |
| Random forest (RF) | 0.197±0.0018 | 0.977±0.0022 | 0.452±0.0055 |
| Elastic net | 0.163±0.0019 | 0.991±0.0025 | 0.427±0.0053 |
| PCA+RF | 0.135±0.0014 | 1.024±0.0023 | 0.404±0.0059 |
| moCluster+RF | 0.107±0.0012 | 1.042±0.0023 | 0.363±0.0060 |
| mixOmics | -0.006±0.0006 | 1.110±0.0016 | 0.110±0.0066 |

**Supplementary Table 2.** Benchmarking six methods to predict drug responses by reporting mean cross-validation performance with 95% confidence interval (CI).

| | $R^2$ mean ± 95%CI | MAE mean ± 95%CI | Pearson's r mean ± 95%CI |
|---|---|---|---|
| **CLP > CCLE** | | | |
| **DeePathNet** | **0.208±0.0118** | **0.933±0.0274** | **0.476±0.0119** |
| Random forest | 0.027±0.0200 | 1.07±0.0315 | 0.390±0.0143 |
| **$CLP^+$ > $CCLE^+$** | | | |
| **DeePathNet** | **0.233±0.0126** | **0.899±0.0262** | **0.532±0.0139** |
| Random forest | -0.106±0.0441 | 1.107±0.0330 | 0.388±0.0183 |

**Supplementary Table 3.** Comparing two methods by evaluating mean generalisation errors with 95% CI of drug response prediction.

| | Accuracy mean ± 95%CI | Macro-average F1-score mean ± 95%CI | AUROC mean ± 95%CI | Stability |
|---|---|---|---|---|
| **DeePathNet** | **0.963±0.0015** | **0.935±0.0030** | **0.998±0.0001** | **0.004** |
| Random forest (RF) | 0.951±0.0018 | 0.895±0.0038 | 0.997±0.0003 | 0.005 |
| *k*-NN | 0.940±0.0018 | 0.894±0.0045 | 0.982±0.0016 | 0.007 |
| PCA+RF | 0.937±0.0023 | 0.885±0.0039 | 0.996±0.0003 | 0.005 |
| moCluster | 0.866±0.0031 | 0.734±0.0034 | 0.987±0.0010 | 0.006 |
| mixOmics+RF | 0.764±0.0040 | 0.881±0.0091 | 0.919±0.0041 | 0.015 |

**Supplementary Table 4.** Benchmarking six methods to predict cancer types by reporting cross-validation performance.

| | Accuracy mean ± 95%CI | Macro-average F1-score mean ± 95%CI | AUROC mean ± 95%CI | Stability |
|---|---|---|---|---|
| **DeePathNet** | **0.902±0.0081** | **0.868±0.0115** | **0.980±0.0030** | **0.019** |
| Random forest (RF) | 0.844±0.0095 | 0.672±0.0178 | 0.969±0.0038 | 0.026 |
| *k*-NN | 0.816±0.0097 | 0.697±0.0200 | 0.897±0.0094 | 0.033 |
| PCA+RF | 0.696±0.0109 | 0.429±0.0111 | 0.880±0.0094 | 0.027 |
| mixOmics | 0.749±0.0525 | 0.676±0.0369 | 0.731±0.0169 | 0.090 |
| moCluster+RF | 0.751±0.0090 | 0.492±0.0152 | 0.917±0.0051 | 0.025 |

**Supplementary Table 5.** Benchmarking six methods to predict breast cancer subtypes by reporting cross-validation performance.

| | Accuracy | Macro-average F1-score | AUROC |
|---|---|---|---|
| **DeePathNet** | **0.902** | **0.798** | **0.971** |
| Random forest | 0.295 | 0.263 | 0.694 |

**Supplementary Table 6.** Comparing two methods by evaluating generalisation errors of breast cancer subtype prediction.
